# Azoospermia and meiotic defects in class I crossover*-*deficient mice are not rescued by increasing an alternate crossover pathway

**DOI:** 10.64898/2026.11.23.768476

**Authors:** Stevan Novakovic, Elissah Granger, Wayne Crismani

## Abstract

Meiotic crossovers are essential for accurate chromosome segregation and fertility. Most crossovers are formed through the so-called class I pathway, which relies on evolutionarily conserved proteins that are collectively referred to as ZMM. In humans, biallelic pathogenic variants in the *ZMM* are associated with non-obstructive azoospermia and premature ovarian insufficiency. *Spo16* is one of the *ZMM* and loss of function causes failure of class I crossover designation, defective synapsis, meiotic arrest, and infertility in mice. On the other hand, increasing crossover rates does not necessarily affect fertility. We hypothesised therefore that increasing crossover rates could rescue some of the reproductive defects of *Spo16*-deficient mice, for which there is precedent in other species. To increase crossover rates, we crossed a novel *Spo16-*deficient mouse strain with the hyper-recombinant *Fancm-*deficient mice*. Fancm* deficiency did not rescue the spermatogenic failure caused by *Spo16* loss. Meiotic arrest, synapsis defects and complete absence of crossover designation remained unaltered in double-mutant males, indistinguishable from *Spo16* deficiency alone. A number of possible explanations exist for our observations, and these are discussed through the lens of mammalian reproductive health.

## Introduction

Meiosis is a specialised cell division in which diploid progenitor cells produce haploid gametes. Meiosis involves a single round of DNA replication followed by two successive cell divisions, reducing the chromosome number from diploid to haploid. Accurate synapsis, crossover formation, and segregation of homologous chromosomes are essential for maintaining the same chromosome number and composition across generations. A defining feature of the first meiotic division is the physical large-scale exchanges between homologous chromosomes through crossovers that are essential for accurate homologue segregation at meiosis I [1,2]. At least one crossover per chromosome pair is required to form bivalents and failure to achieve this produces univalents, anaphase I segregation errors, aneuploid gametes, and infertility [1,3]. Total crossover numbers and crossover distribution along individual chromosomes are under tight regulatory control, with mechanisms operating at the level of both crossover designation and spatial patterning [2]. Another defining feature of meiosis I in most species studied is the tripartite synaptonemal complex which is composed of the chromosomal axes of two homologues, and the central element. In mammals the axes can be visualised by labelling the protein SYCP3, and the central element by labelling SYCP1 [4,5].

Meiotic recombination is initiated by programmed DNA double-strand breaks (DSB) generated by SPO11 and accessory proteins [6–8]. Processed DSB ends invade the homologous chromosome to form displacement loops, which can be resolved as either crossover or non-crossover products, with pathway choice established early in the recombination program [9,10]. In mammals, and most eukaryotes [1,11], the majority of crossovers form through the class I pathway, which depends on the MutL heterodimer MLH1-MLH3 together with a conserved group of factors collectively known as the ZMM proteins, including SYCP1/ZIP1, SHOC1/ZIP2, RNF212/ZIP3, ZIP4/TEX11, MER3/HFM1, MSH4-MSH5, and SPO16 [12,13]. Within the ZMM, SPO16 forms a sub-complex with SHOC1 and TEX11 (collectively term the conserved ZZS complex) and is required for class I crossover formation [14,15].

Crossovers which arise through the class I pathway are more evenly spaced along chromosomes than would occur by chance, and this phenomenon is known as crossover interference [16]. Loss of ZMM components typically causes defective synapsis, failure to form crossovers, meiotic arrest at zygotene or pachytene, and infertility. A minority of meiotic crossovers arise through a class II pathway, mediated principally by the structure-selective endonuclease MUS81-EME1, which are not sensitive to interference [17–20].

The non-redundant requirement of ZMM-dependent crossover formation in the mammalian germline has been demonstrated in human and broader mammalian genetics. Loss-of-function variants in TEX11 are found in men with non-obstructive azoospermia and meiotic arrest [21]; biallelic SHOC1 mutations cause azoospermia with pachytene arrest [22]; *Spo16*-deficient mice display defective synapsis, failure to form normal crossover sites, meiotic arrest and complete infertility [14,15]; MSH4 and MSH5 variants are associated with primary ovarian insufficiency and have also been identified in male infertility cohorts [23–27]. SYCP1 variants have been linked to severe spermatogenic failure including oligozoospermia [28]. Collectively, these observations support the view that disruption of class I crossover and synapsis factors is poorly tolerated in the mammalian germline.

In parallel, FANCM serves as a conserved anti-crossover factor [29–34]. FANCM is an ATP-dependent DNA translocase with established *in vitro* roles in the remodelling of branched DNA structures [35–38]. In meiosis, FANCM is thought to disassemble recombination intermediates and direct DSB repair toward non-crossover outcomes, thereby acting as a negative regulator of crossover formation [29,39].

In *Arabidopsis thaliana*, loss of *fancm* substantially increases meiotic crossover frequency in inbred lines. In ZMM-defective backgrounds, *fancm* restores bivalent formation in *zip4*, *shoc1*, *msh5* and *msh4* mutants; *fancm* alleles were additionally recovered as fertility suppressors in an *hei10* screen [29,40,41]. Therefore, we sought to test the hypothesis that loss of function of *Fancm* in a *Spo16-*deficient mouse could rescue the meiotic defects and azoospermia.

An anti-crossover role for FANCM has been observed in mammals, where *Fancm* loss increases meiotic crossover frequency [33,42] which provided a key tool to test this question of azoospermia rescue. We generated *Spo16 Fancm* double-mutant mice on a C57BL/6 background and assessed fertility, spermatogenesis, meiotic progression, and synapsis across genotypic combinations. *Fancm* loss did not rescue the *Spo16-*deficient meiotic phenotypes. Double-mutant males retained the severe spermatogenic failure and meiotic disruption seen in *Spo16* single mutants, indicating that increased crossovers are not sufficient to compensate for loss of *Spo16*-dependent crossover formation in mammals.

## Methods

### Ethics statement and animals

All animal procedures were approved by the St. Vincent’s Hospital Melbourne Animal Ethics Committee under approval numbers 002/20-r1 and 014/23-r1 and conducted in accordance the Australian National Health and Medical Research Council Act 1992 and the Australian code for the care and use of animals for scientific purposes (2013). Mice were maintained on a C57BL/6 background under specific pathogen-free conditions, with food and water provided *ad libitum*.

### Mouse production, breeding and genotyping

The *Spo16* mutant mouse line was generated by SA Genome Editing (SAGE; Adelaide, Australia) using CRISPR-Cas9 genome editing on a C57BL/6J background. Two guide RNAs were used to target the *Spo16* locus, with guides positioned in introns 2 and 4 to generate a deletion spanning exons 3 and 4: CATTGAGATATCGGATCAAG and GTTATATGTAACCCAACAGG. Transmitted mutant alleles were confirmed by Sanger sequencing and segregation analysis before more detailed characterisation.

The *Fancm* loss-of-function line has been described previously and was maintained on a C57BL/6 background [33]. *Spo16* and *Fancm* mutant alleles were combined through heterozygous carrier crosses to generate *Fancm^+/^*^−^*;Spo16^+/^*^−^ double-heterozygous animals. Experimental animals were generated from heterozygous intercrosses, and males from the following four genotypes were analysed: *Fancm^+/+^;Spo16^+/+^,Fancm^−/-^;Spo16^+/+^, Fancm^+/+^;Spo16^−/-^* and *Fancm^−/-^;Spo16^−/-^*.

Genotyping was performed by PCR using primers listed in Supplementary Table S1.

### Reproductive phenotyping

Adult males aged 6 to 9 weeks were weighed before testes were dissected and weighed individually. For each male, mean testis weight was calculated as the average of the left and right testes. Relative testis weight was calculated by normalising mean testis weight to body weight. Testes were then allocated for histological analysis, daily sperm production, protein extraction or meiotic chromosome spreads as required.

For histological analysis, testes were fixed in Bouin’s solution overnight, alcohol processed, embedded in paraffin wax, and sectioned using standard methods. Dewaxed testis sections were stained with periodic acid-Schiff reagent and haematoxylin, dehydrated, and mounted under coverslips with DPX. Sections were used to assess seminiferous tubule architecture, germ cell composition and the presence or absence of mature sperm. Brightfield images of testis sections were acquired using a 40x air objective on an Olympus IX81 microscope with an Olympus DP80 camera and cellSens Dimension software (v4.3.1, Build 32730), and processed in Fiji (ImageJ v1.53).

Daily sperm production was determined from snap-frozen testes using a Triton X-100 nuclear solubilisation method, as described previously [33]. Briefly, testes were homogenised in DSP buffer containing 0.15 M NaCl, 0.01% NaN_3_ and 0.05% Triton X-100 to lyse somatic cells and remove residual sperm tails while retaining condensed spermatid nuclei. Homogenates were counted using a Neubauer haemocytometer, and daily sperm production per testis was calculated by dividing the number of elongated spermatids per testis by 4.84, corresponding to the estimated number of days elongated spermatids remain in the mouse testis.

### Cytological analysis of meiotic prophase I

Meiotic chromosome surface spreads were prepared from testes of males aged 6 to 8 weeks using a hypotonic-paraformaldehyde spreading method, as previously described [33,43]. Briefly, testes were disrupted in PBS to release meiocytes, which were then exposed to a hypotonic sucrose solution and dispensed onto slides coated with freshly prepared fixative solution containing 1% paraformaldehyde (pH 9.2) and 0.15% Triton X-100. Slides were dried for up to 2 hours in a humid chamber, washed in 0.4% Photo-Flo (Kodak), and air-dried.

For meiotic prophase I staging and synapsis analysis, chromosome surface spreads were stained with primary antibodies against SYCP1 and SYCP3. For MLH3 focus analysis, separate chromosome surface spreads were stained with primary antibodies against MLH3 and SYCP3. Primary antibodies were applied overnight, followed by Alexa Fluor-conjugated secondary antibodies. Slides were counterstained with DAPI and mounted in ProLong Diamond Antifade Mountant (Invitrogen) under a glass coverslip. Antibody sources and dilutions are listed in Supplementary Table S2.

Images were acquired on an Olympus IX81 microscope using a 100x oil-immersion objective, an Olympus DP80 camera, and cellSens Dimension software (v4.3.1, Build 32730), and processed in Fiji (ImageJ v1.53)

Meiotic prophase I staging followed established SYCP1 and SYCP3 criteria. Leptotene nuclei were identified by short, discontinuous SYCP3 axial elements with minimal or absent SYCP1. Early zygotene nuclei displayed elongating but still discontinuous SYCP3 axes with limited regions of SYCP1 marking the initiation of synapsis. Late zygotene nuclei showed extensive but incomplete synapsis, with SYCP1 present on most, but not all, autosomal axes and the XY pair remaining largely unsynapsed. Pachytene nuclei were defined by full synapsis of all 19 autosomal bivalents, with continuous SYCP1 co-localising along SYCP3 axes. Diplotene nuclei were identified by desynapsis and loss of SYCP1 along separating axes.

In *Spo16-*deficient genotypes, nuclei were classified as zygotene-like or pachytene-like using criteria commonly applied to synapsis-defective mutants, as previously described [15,44,45]. Zygotene-like nuclei exhibited extensive axial element elongation, with SYCP3 forming irregular, branched, or partially collapsed structures rather than organised linear axes, and SYCP1 present on multiple chromosome segments in a fragmented, uneven pattern that could not be cleanly assigned to early or late zygotene by autosomal count. Pachytene-like nuclei were defined as *Spo16-*deficient spermatocytes with extensive SYCP1 staining and elongated SYCP3 axes consistent with progression beyond early zygotene, but lacking the fully organised autosomal synapsis seen in wild-type pachytene cells. These nuclei retained abnormal SYCP3 morphology, including tangled, branched or collapsed axial structures, and regions of incomplete or irregular SYCP1/SYCP3 co-localisation.

For meiotic prophase I staging, three slides were analysed per mouse from three mice per genotype. For each slide, the first 100 prophase I nuclei encountered were scored, giving at least 300 nuclei per mouse and at least 900 nuclei per genotype.

### Quantification of synapsis

Quantitative analysis of synapsis was restricted to nuclei classified as pachytene or pachytene-like. For each nucleus, SYCP3 was used to identify chromosome axes and SYCP1 was used as a readout of completed synapsis. Continuous stretches of SYCP1 along chromosome axes were traced using the segmented line tool in Fiji, with clear gaps or branch points used to delimit individual segments. Total SYCP1 length per nucleus was calculated as the sum of all continuous SYCP1-positive segments.

MLH3 foci were counted manually in Fiji on MLH3/SYCP3-stained chromosome surface spreads. MLH3 analysis was restricted to pachytene or pachytene-like nuclei. Discrete MLH3 foci associated with SYCP3-labelled chromosome axes were counted, while diffuse background signal or off-axis staining was excluded. At least 30 pachytene or pachytene-like nuclei were examined per mouse, with at least three mice analysed per genotype.

For SYCP1 length and MLH3 focus analyses, nuclei with overlapping chromosomes, severe folding artefacts, or ambiguous chromosome-axis morphology were excluded.

### Statistics

Statistical analyses were performed using GraphPad Prism (11.0). Genotype frequencies were compared to Mendelian expectations by chi-square test. Differences in body weight, testis weight, relative testis weight and daily sperm production were assessed by one-way ANOVA with Tukey’s multiple comparisons test.

For meiotic prophase I staging, values were calculated per mouse from three slides, with 100 nuclei scored per slide. For SYCP1 length and MLH3 focus analyses, values were first summarised per mouse, and genotype comparisons were performed using mouse-level values as the biological replicate. Individual nuclei are shown in the figures to display within-animal variation, but statistical testing was performed using n = 3 males per genotype. Genotype comparisons for SYCP1 length and MLH3 foci were performed by one-way ANOVA with Tukey’s multiple comparisons test. A value of *P* < 0.05 was considered statistically significant.

## Results

### Generation and initial characterisation of a *Spo16* mutant mouse line

To investigate whether *Fancm* loss modifies the meiotic defects caused by loss of *Spo16*, we generated *Fancm*^−^*^/^*^−^;*Spo16*^−^*^/^*^−^ double-mutant mice on a C57BL/6 background. The *Fancm* null allele used here has been described previously [33] and has been shown to result in complete loss of *Fancm* function.

We generated a *Spo16* loss-of-function allele using CRISPR-Cas9-mediated genome editing. Because complete disruption of *Spo16* was expected to cause sterility and preclude germline transmission, we designed a dual-guide strategy to facilitate recovery of fertile heterozygous founders. Guide RNAs were positioned within introns 2 and 4, flanking exons 3 and 4. Simultaneous cleavage at both sites on one chromosome was expected to generate a readily detectable deletion encompassing these exons, whereas repair events affecting only one guide site on the second chromosome would be confined to intronic sequences and were therefore less likely to disrupt *Spo16* function.

We use the exon-numbering convention of Zhang *et al.* (2019), in which the *Spo16* coding sequence extends from exons 2 to exon 6 [15]. Under this convention, deletion of exons 3 and 4 was expected to disrupt the coding sequence. Consistent with this, Zhang *et al.* showed that a 19-bp insertion within exon 3 caused a frameshift and premature termination codon, resulting in loss of SPO16 function.

A total of 187 zygotes were injected with both guide RNAs, yielding 13 founders, three of which carried candidate deletion alleles by PCR screening. Founder #11 transmitted a mutant allele to its progeny. Sequencing of the transmitted allele identified an 8-bp deletion at the 5′ guide site and a 3,255-bp deletion encompassing exons 3 and 4. This allele is predicted to shift the reading frame and introduce a premature termination codon early in exon 5 (Figure 1A).

**Figure 1.**
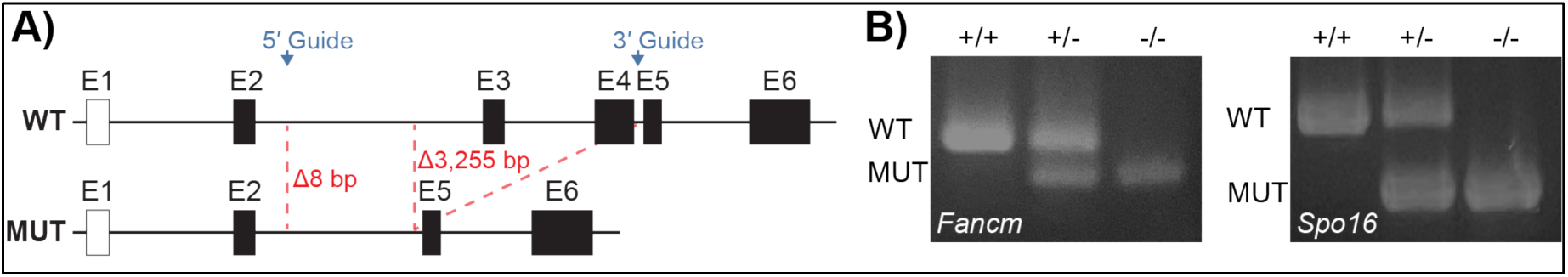
Generation and validation of *Spo16* and *Fancm* mutant mouse lines. A) Schematic of the CRISPR-Cas9-generated *Spo16* founder 11 mutant allele. WT and MUT tracks show the wild-type and mutant alleles, respectively. Exons are numbered according to the six-exon *Spo16* transcript model, including non-coding exon 1 (open box) and coding exons (filled boxes). The mutant allele contains an 8 bp deletion at the 5′ guide site and a 3,255 bp deletion spanning exons 3-4, resulting in a frameshift and premature stop codon in exon 5. Intronic distances are compressed for clarity. B) Representative genotyping PCR distinguishing wild-type, heterozygous and homozygous mutant alleles for *Fancm and Spo16*.

Colonies were maintained as *Fancm^+/^*^−^*;Spo16^+/^*^−^ double heterozygotes. PCR genotyping readily distinguished wild-type, heterozygous, and homozygous mutant alleles at both loci (Figure 1B). The observed distribution of offspring genotypes is shown in Supplementary Table S3. At the *Spo16* locus, genotype frequencies were consistent with Mendelian expectations (*n* = 362; 77 *Spo16*^+/+^, 197 *Spo16*^+/−^, and 88 *Spo16*^−/−^; χ² = 3.50, df = 2, *P* = 0.174). In contrast, *Fancm*^−/−^ offspring were underrepresented (99 *Fancm*^+/+^, 202 *Fancm*^+/−^, and 61 *Fancm*^−/−^; χ² = 12.85, df = 2, *P* = 0.0016), consistent with previous observations [33,46,47].

We next assessed body and testis weights as an initial measure of reproductive phenotype. Loss of *Fancm*, but not *Spo16*, reduced body weight (two-way ANOVA: Fancm, F(1,32) = 11.09, P = 0.002; Spo16, F(1,32) = 0.41, P = 0.52; interaction, F(1,32) = 0.56, P = 0.46). Testis weight was reduced in all mutant groups relative to wild type (Figure 2B). *Fancm*^−/−^ males showed an approximately 54% reduction in mean testis weight (37.3 ± 5.4 mg vs 81.2 ± 11.9 mg, *P* < 0.0001, two-way ANOVA on log-transformed data), consistent with prior characterisation of this allele on the C57BL/6 background [33]. *Spo16*^−/−^ males showed a more severe reduction in testis weight (23.6 ± 2.4 mg, *P* < 0.0001). Double-mutant males had the lowest absolute testis weight of any group (14.3 ± 1.9 mg, *P* <0.0001, two-way ANOVA on log-transformed data), significantly lower than *Spo16^−/−^*single mutants (p<0.0001). After normalisation to body weight however, relative testis weight all genotype comparisons were significant (two-way ANOVA: Fancm, F(1,32) = 156.0, P < 0.0001; Spo16, F(1,32) = 538.8, P < 0.0001) except for between *Spo16^−/−^* and double-mutant males after Tukey post-hoc correction (1.11 ± 0.11 vs 0.91 ± 0.33 mg/g, *P* = 0.33, Figure 2C).

**Figure 2.**
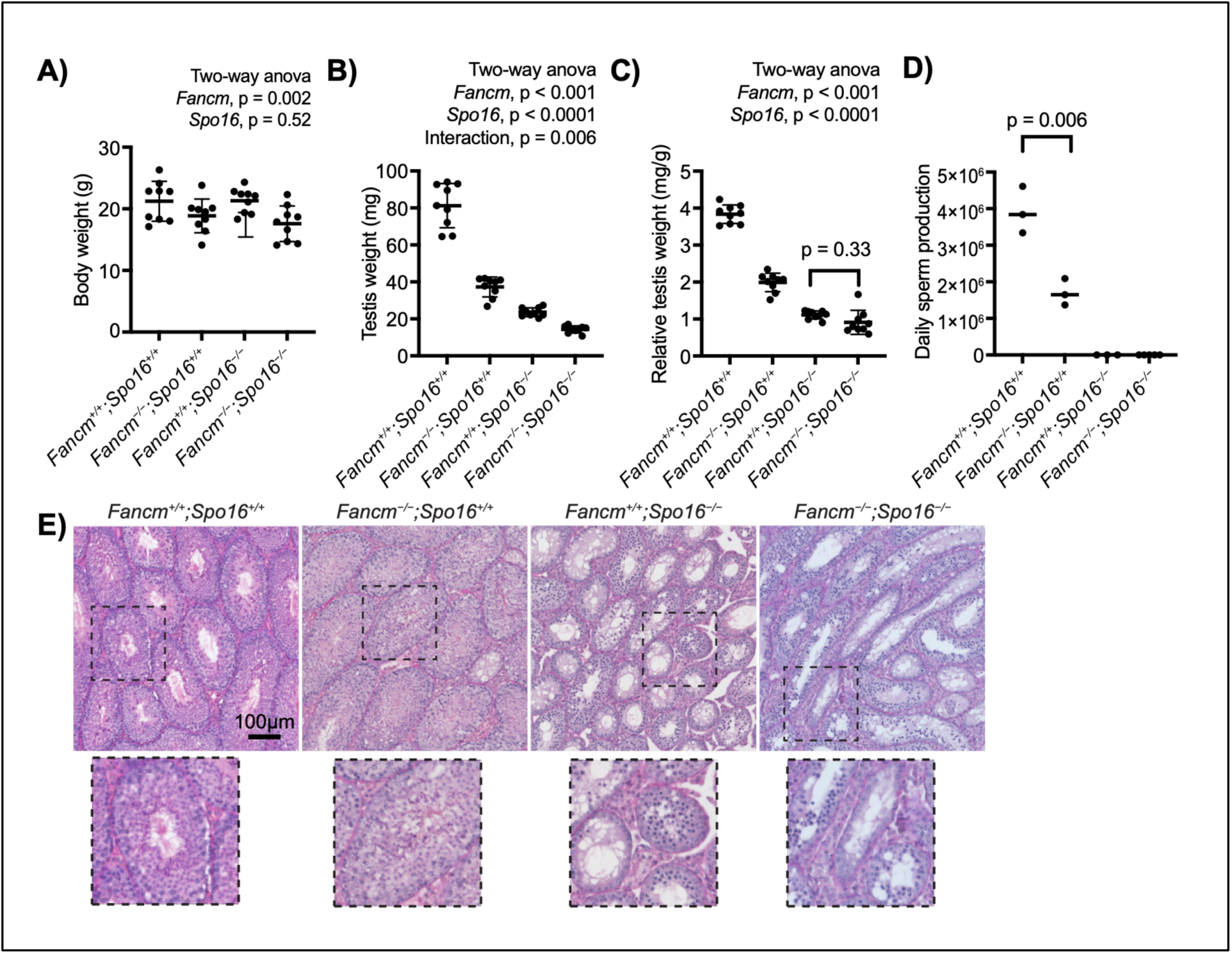
*Fancm* loss does not restore testis weight or sperm production in *Spo16*-deficient males. A) Body weights. Loss of *Fancm*, but not *Spo16*, reduced body weight (two-way ANOVA: Fancm, F(1,32) = 11.09, P = 0.002; Spo16, F(1,32) = 0.41, P = 0.52; interaction, F(1,32) = 0.56, P = 0.46). B) Mean testis weight was reduced by loss of function of *Fancm* or *Spo16*, and through their interaction in the double knockout. *Fancm*, F(1,32) = 190.1, P < 0.0001; *Spo16*, F(1,32) = 551.4, *P* < 0.0001), *Fancm Spo16* (F(1,32) = 8.75, P = 0.006). C) After normalisation to body weight, relative testis weight all genotype comparisons were significant (two-way ANOVA: *Fancm*, F(1,32) = 156.0, P < 0.0001; *Spo16*, F(1,32) = 538.8, P < 0.0001) except for between *Spo16^−/−^* and double-mutant males after Tukey post-hoc correction (1.11 ± 0.11 vs 0.91 ± 0.33 mg/g, *P* = 0.33. D) Azoospermia in *Fancm^+/+^;Spo16^−/−^*was not rescued in *Fancm^−/−^;Spo16^−/−^*. E) Representative PAS-stained testis sections from each genotype. *Fancm^+/+^;Spo16^+/+^*, *Fancm^−/−^;Spo16^+/+^* testis contained seminiferous tubules with mature sperm, whereas *Fancm^+/+^;Spo16^−/−^*, and *Fancm^−/−^;Spo16^−/−^* testes contained spermatocytes but lacked mature sperm. Dashed boxes indicate regions shown at higher magnification below. Scale bar, 100 µm. Data are individual mice with mean ± SD. For A - C, two-way ANOVA with Tukey’s post-hoc correction was used. For panel D, an unpaired t-test was used for genotypes where non-zero values were obtained.

### *Fancm* loss does not rescue the spermatogenic failure observed in S*po16-*deficient males

*Spo16* is required for class I crossover formation, and *Spo16* loss causes complete spermatogenic failure. We therefore asked whether *Fancm* deficiency, which increases meiotic crossover frequency, could compensate for *Spo16* loss. We next measured daily sperm production and examined testis histology.

Daily sperm production confirmed severe spermatogenic failure in *Spo16*-deficient males (Figure 2D). *Fancm^−/−^* males had reduced but detectable sperm output relative to wild-type males (1.70 ± 0.36 × 10⁶ versus 3.93 ± 0.64 × 10⁶ sperm per day, unpaired t-test, *P* = 0.006), consistent with previous findings. Sperm production was undetectable in both *Fancm^+/+^;Spo16^−/−^* and *Fancm^−/−^;Spo16^−/−^* males. Histological analysis supported the absence of post-meiotic germ cells, with no detectable immature or mature sperm in seminiferous tubules from either *Spo16*-deficient genotype. Spermatogonia and cells morphologically consistent with pachytene-arrested spermatocytes were nevertheless present (Figure 2E). Therefore, loss of function of *Fancm* loss does not restore post-meiotic germ-cell development or sperm production in *Spo16-*deficient males.

### Meiotic progression and synapsis defects in *Spo16-*deficient male mice are not rescued by loss of *Fancm*

Synapsis and crossover designation are central features of meiotic prophase I progression and linked to class I crossover formation [1,12,13]. To assess whether *Fancm* deficiency modifies meiotic prophase I progression in the absence of *Spo16*, we analysed chromosome synapsis and crossover designation by immunofluorescence staining of SYCP1 and SYCP3 on chromosome surface spreads from adult seminiferous tubules.

In wild-type and *Fancm^−/−^* spermatocytes, meiotic prophase I proceeded normally, with cells distributed across leptotene, zygotene, pachytene and diplotene stages in comparable proportions, consistent with previous observations [33] (Figure 3B). Approximately half of all scored spermatocytes were at diplotene in both genotypes (wild type: 50.8 ± 2.7%; *Fancm^−/−^*: 48.7 ± 2.5%), which is consistent with normal meiotic progression. In contrast, both *Spo16^−/−^* and *Fancm^−/−^;Spo16^−/−^* spermatocytes showed a marked redistribution of prophase I stages. Leptotene and zygotene-like cells were elevated relative to controls, early zygotene accounted for approximately 28-29% of scored spermatocytes in both *Spo16*-deficient genotypes compared with approximately 5% in controls. Diplotene cells were entirely absent from both *Spo16^−/−^*and double-mutant testes (Figure 3B), indicating failure of the meiotic program to progress past the pachytene. The staging profiles of *Spo16^−/−^* and *Fancm^−/−^;Spo16^−/−^* spermatocytes were indistinguishable, with cells accumulating at zygotene-like and pachytene-like stages as previously described for *Spo16*-deficient spermatocytes [15].

**Figure 3.**
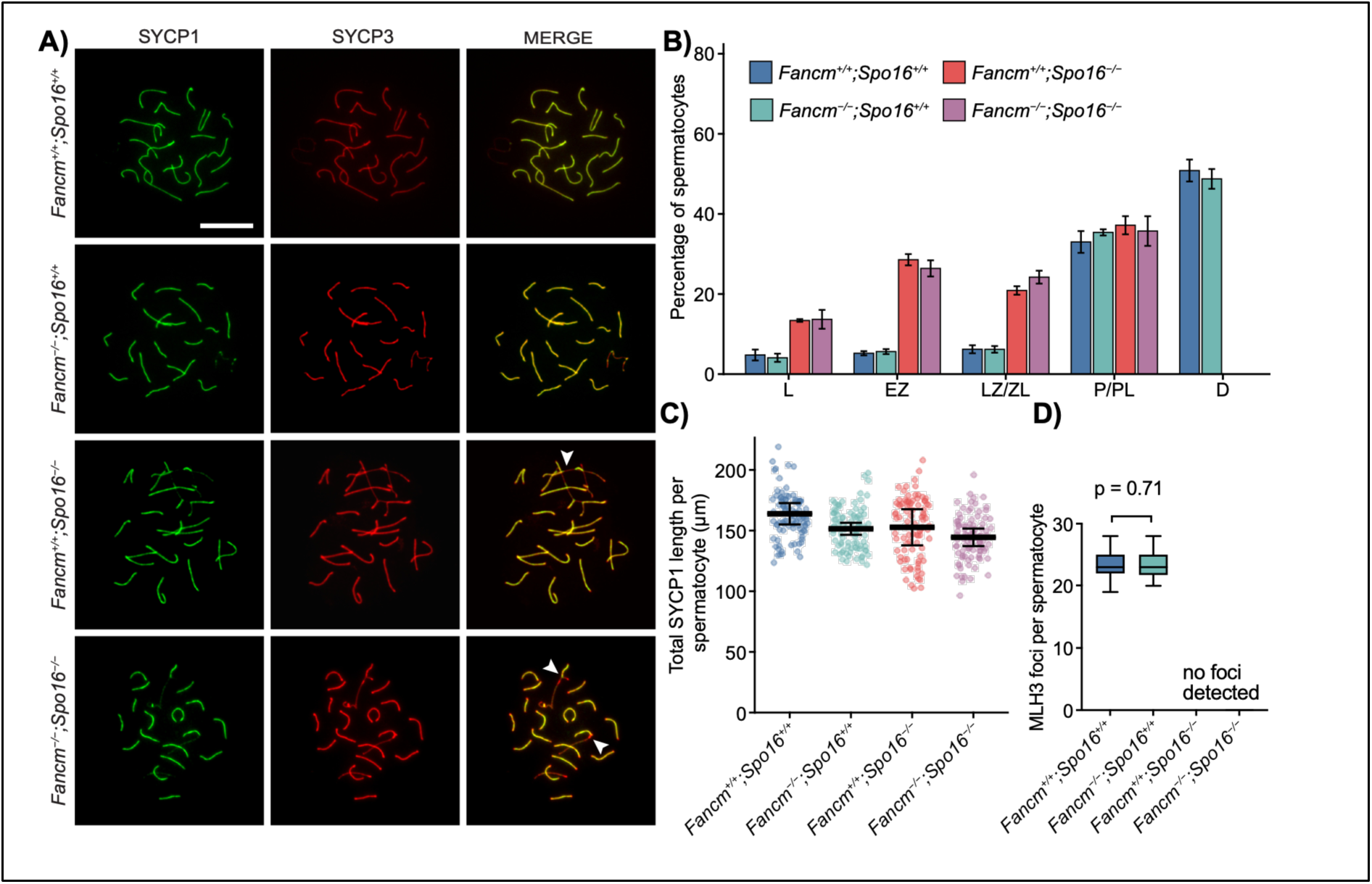
*Fancm* loss does not restore meiotic progression or synapsis in *Spo16*-deficient spermatocytes. A) Representative nuclear surface spreads of spermatocytes stained for SYCP1 and SYCP3. *Fancm^+/+^;Spo16^+/+^ and Fancm^−/−^;Spo16^+/+^* cells are shown at pachynema, while *Spo16*-deficient cells are shown at a pachynema-like stage. White arrowheads indicate abnormal SYCP1/SYCP3 organisation in *Spo16*-deficient spermatocytes. Scale bar, 20 µm. B) Meiotic prophase I staging. Bars show mean percentage of spermatocytes per mouse ± SD from n = 3 males per genotype. L, leptotene; EZ, early zygotene; LZ/ZL, late zygotene or zygotene-like; P/PL, pachytene or pachytene-like; D, diplotene. C) Total SYCP1 length per spermatocyte. Points represent individual spermatocytes; statistical comparisons were performed using per-mouse means. Bars show mean ± SD; two-way ANOVA, *Spo16*, *P* = 0.14; *Fancm*, *P* = 0.10, n = 3 mice per genotype). D) Quantification of MLH3 foci per spermatocyte. MLH3 foci were detected in *Fancm^+/+^;Spo16^+/+^*and *Fancm^−/−^;Spo16^+/+^* pachytene spermatocytes but were absent from *Spo16-*deficient pachytene-like spermatocytes. At least 30 spermatocytes from each of n = 3 males per genotype were quantified. ns, not significant; with an unpaired t-test on per-mouse means *P* = 0.71).

Consistent with the staging data, SYCP1 staining revealed abnormal synapsis in both *Spo16*-deficient genotypes. Wild-type and *Fancm^−/−^* pachytene spermatocytes displayed organised, continuous SYCP1 along paired autosomal axes, whereas *Spo16^−/−^*and *Fancm^−/−^;Spo16^−/−^* pachytene-like spermatocytes showed abnormal SYCP1 organisation, including fragmented and irregular staining associated with abnormal axial structures (Figure 3A, arrows). Quantification of total SYCP1 length per spermatocyte revealed a significant, albeit modest 1.4 µm reduction, in SYCP1 length per filament (*Spo16*, F(1,8) = 25.5, *P* = 0.001; *Fancm*, F(1,8) = 4.81, *P* = 0.060; interaction (*Fancm^−/−^;Spo16^−/−^*), F(1,8) = 0.02, *P* = 0.89). The number of filaments per nucleus also increased modestly in *Spo16-*deficient mice compared to the wildtype, consistent with incomplete synapsis (19.0 vs 21.2; F(1,8) = 71.1, P < 0.0001). The defects in *Spo16*-defecient mice are broadly similar to what has been described previously [15]. Loss of *Fancm* did not alter mean filament lengths (F(1,8) = 4.81, *P* = 0.060) or filament number per nucleus (*P =* 0.33), similar to previous observations, and there was no interaction between the two genes (*P* = 0.89 and *P* = 0.32). The total SYCP1 length per nucleus was not different between genotypes and no interaction was detected (*Spo16*, *P* = 0.14; *Fancm, P* = 0.10, n = 3 mice per genotype).

MLH3, a marker of class I crossover designation, was detected at a mean of 23.1 ± 2.1 foci per pachytene spermatocyte in wild-type males and 23.2 ± 2.1 foci per *Fancm^−/−^* males, with no significant differences between these genotypes (*P* = 0.71, Figure 3D). MLH3 foci were entirely absent from pachytene-like spermatocytes in both *Spo16^−/−^*and *Fancm^−/−^;Spo16^−/−^* males. Loss of *Fancm* therefore does not restore class I crossover designation in *Spo16*-deficient spermatocytes, which would be consistent with our previous work [33].

Taken together, these data demonstrate that *Fancm* deficiency does not rescue the meiotic prophase I defects caused by loss of *Spo16*. The staging arrest, synapsis abnormalities, and absence of MLH3 foci observed in *Spo16^−/−^* spermatocytes is also seen in *Fancm^−/−^;Spo16^−/−^*double mutants, therefore indicating that increased class II crossover potential is not sufficient to compensate for the loss of *Spo16*-dependent recombination in mammals.

## Discussion

Here, we generated a novel loss-of-function *Spo16* allele. Homozygotes recapitulated key features of *Spo16* deficiency previously described, including severe spermatogenic failure, absence of post-meiotic cells, abnormal synapsis, prophase I arrest, and loss of MLH3 foci [15]. We then used this azoospermic model to test whether increasing crossover activity through loss of *Fancm* could compensate for the failure of class I crossover formation. Across all phenotypic measures examined, including testis weight, sperm production, seminiferous tubule histology, meiotic staging, synapsis, and MLH3 focus formation, *Fancm*^−/−^;*Spo16*^−/−^ males showed no evidence of rescue relative to *Spo16*^−/−^ males. Similarly, loss of *Fancm* did not restore meiotic progression or spermatogenesis in *Spo16*-deficient males. Therefore, the clinical implication of this work is that increasing class II crossovers, does not appear to provide a therapeutic route to severe infertility, caused by complete loss of function of the class I crossover pathway.

A first question we sought to address here was whether the known increase in crossover frequency caused by *Fancm* loss in mice was sufficient to compensate for complete *Spo16* deficiency. Available genetic mapping and single-gamete crossover data show that *Fancm* loss produces a hyper-recombination phenotype in otherwise recombination-competent mice [33,42]. However, *Spo16*-deficient spermatocytes lack the ZZS-dependent class I crossover programme and show defective synapsis and prophase I arrest [15]. Thus, based on the established magnitude of the *Spo16-*deficient phenotype in mice, loss of *Fancm* might not be expected to replace the extensive class I crossover failure caused by complete loss of *Spo16*.

Nevertheless, the *Spo16*-deficient background was selected to provide a sensitive test for cytogenetic rescue that *Fancm* could have potentially conferred. In *Arabidopsis*, the *fancm* hyper-recombination phenotype depends strongly on genetic background: *fancm* restores bivalent formation in several ZMM-mutant inbred backgrounds, whereas this dramatic increase in crossover frequency was reduced in F1 hybrids [29,32,41,48]. Similarly, we previously observed in F1 *Fancm-*deficient mice in a cross between FVB and C57BL/6 strains, that the increase in crossover frequencies compared to their wild-type counterparts was modest compared to the levels of increase seen in *Arabidopsis*. Therefore, the *Spo16* strain was selected not only to provide a sensitive context in which to detect cytogenetic rescue, but also to enable assessment of *Fancm*-dependent crossover effects in an inbred C57BL/6 background, where these potential hyper-recombination phenotypes were hypothesised to be maximal. Because an inbred background lacks informative polymorphisms for genetic mapping [49], crossover events were quantified cytogenetically. However, any additional crossover activity sufficient to improve meiotic function should have produced detectable improvement in synapsis, seminiferous tubule histology, meiotic progression, or histological detection of post-meiotic spermatogenesis stages. No such phenotypic rescue was observed.

Another possibility for why no phenotypic rescue was observed may be linked to the strength of the pachytene checkpoint that is triggered by a severe reduction of class I crossovers in mice. *Spo16*-deficient spermatocytes display extensive abnormal synapsis and fail to progress beyond a pachytene-like stage as seen here and previously [15]. Therefore, it is possible that even a hypothetical large boost of crossovers through the class II pathway may not be sufficient to rescue meiotic progression. In male meiosis, persistent autosomal asynapsis recruits ATR and BRCA1 to unsynapsed chromosome axes and can impair meiotic sex chromosome inactivation by preventing their appropriate concentration on the XY bivalent [50,51]. Zhang *et al.* showed that *Spo16*-deficient spermatocytes form a pseudosex body in which γH2AX and BRCA1 are distributed across unsynapsed autosomal chromatin, accompanied by incomplete silencing of *Zfy1* and *Zfy2* [15]. Thus, meiotic sex chromosome inactivation is already compromised in these *Spo16-*deficient meiocytes.

Other anti-crossover pathways could, in principle, have a greater capacity to alter crossover output than loss of *Fancm* alone. In plants, the BLM homologue RECQ4, TOP3α and RMI1 act in parallel to FANCM to limit crossovers [30,52], while FIGNL1 acts through a distinct anti-crossover mechanism [41]. However, FANCM itself has functions beyond crossover control, including in germ-cell maintenance and genome stability and may cause other cell cycle phenotypes, as reflected by reduced spermatogenesis and fertility [33,47]. Similarly, all genes from the Fanconi anaemia DNA repair pathway are required for efficient gametogenesis and normal levels of fertility [53]. More strikingly, germline-specific loss of FIGNL1 or FIRRM/FLIP in mice causes defective meiotic DSB repair, failure of homologous synapsis, prophase I arrest, and apoptosis [54] rather than a relatively benign increase in crossover numbers as was observed in plants [41]. Thus, although a number of key features of crossover control are conserved between plants, mammals, yeasts and eukaryotes more broadly, perturbation of individual anti-crossover pathways can have substantially different biological consequences.

Together, these findings may define a boundary on the ability of anti-crossover manipulation to compensate for meiotic failure. In the context of complete *Spo16* loss, *Fancm* deficiency does not restore synapsis, meiotic progression, or spermatogenesis. This does not exclude the possibility that some combination of loss, of anti-crossover factors may compensate for loss of function or reduced activity of a pro-crossover factor in the context of meiosis. For example, *Blm* represents an alternative target to *Fancm* which may lead to an increase in crossovers, however a hypomorphic allele would be required given the lethality of complete loss of function. Similarly hypomorphic alleles of ZMMs such as *Spo16*, *Shoc1*, or *Tex11* that retain some residual activity of recombination-intermediate maturation, or synapsis might provide more informative models in which to test this possibility, however these are beyond the scope of this study.

## Acknowledgements

The *Spo16* mutant mouse line was generated by Mel White and Professor Paul Thomas of SA Genome Editing (SAGE; the University of Adelaide, Australia) and the Australian Phenomics Network.

WC received fellowships and funding related to this work from the Australian National Health and Medical Research Council (GNT1129757, GNT1185387). Thank you to Arnaud DeMuyt and Valerie Borde for helpful discussions.

## Authors’ contributions

All authors wrote, reviewed and discussed the manuscript. SN prepared the figures. All authors read and approved the final manuscript.

## upplementary Material

**Supplementary Table S1.**
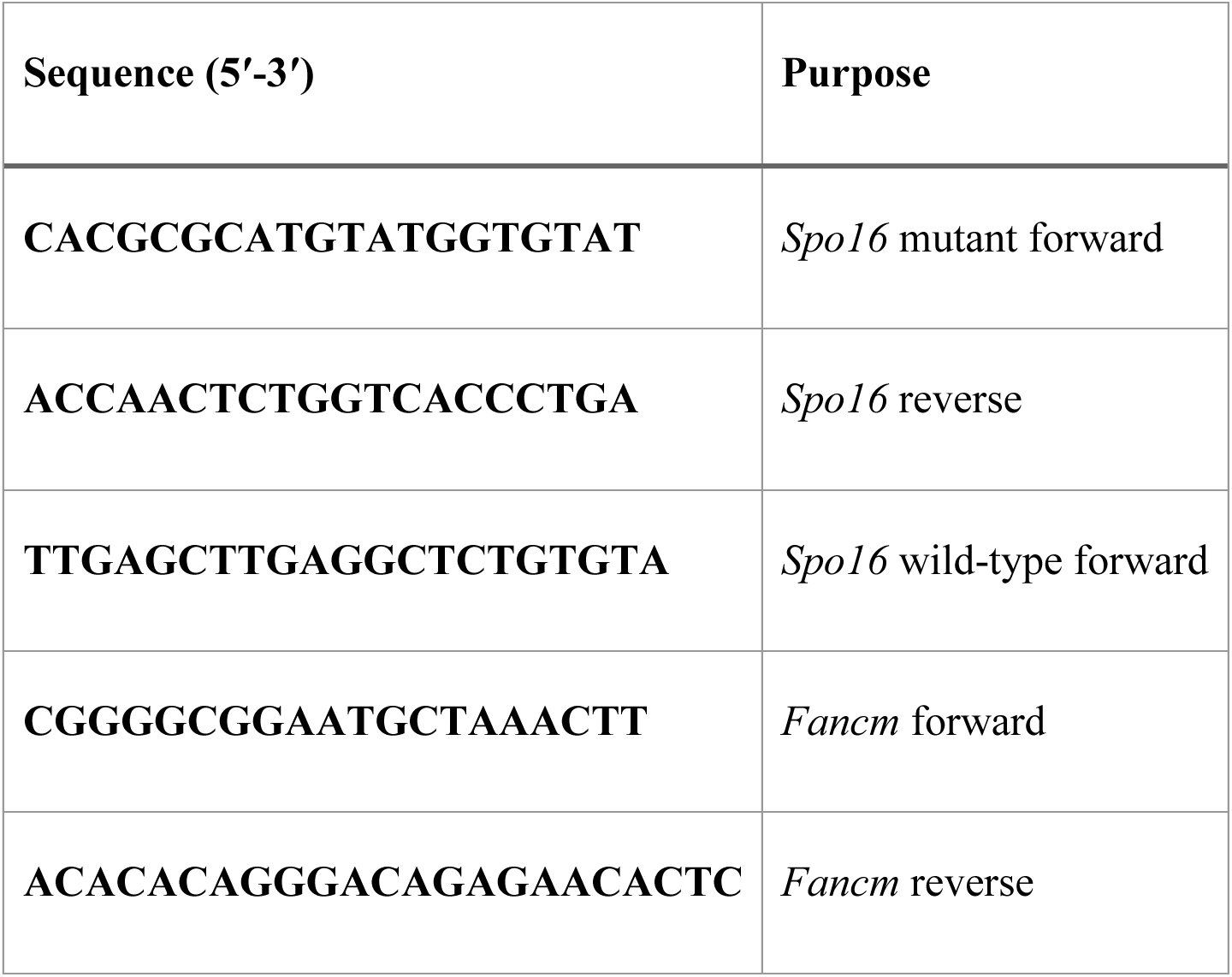
Genotyping primers.

**Supplementary Table S2.**
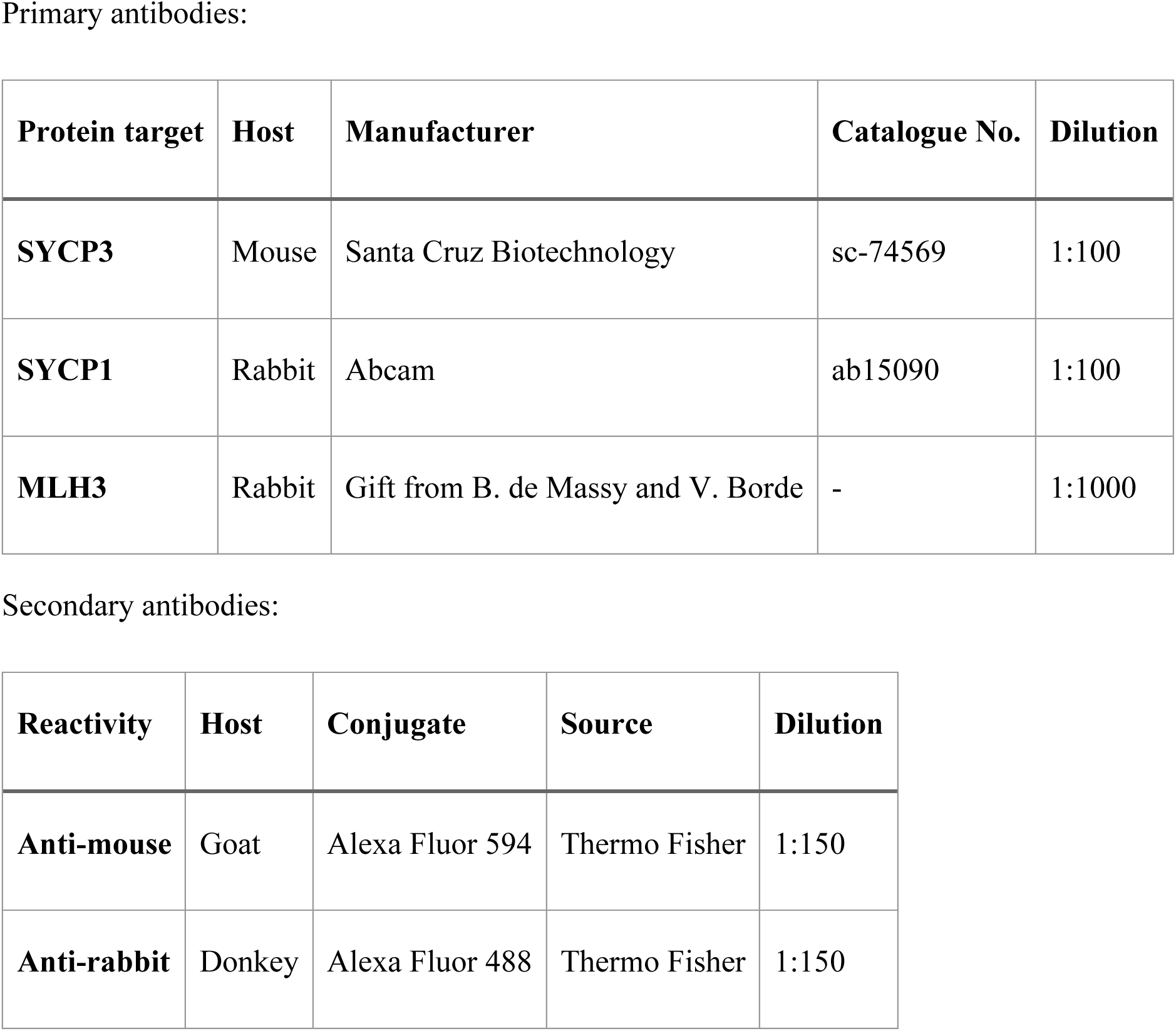
Antibodies.

| Primary antibodies: |  |  |  |  |
|---|---|---|---|---|
| **Protein target** | **Host** | **Manufacturer** | **Catalogue No.** | **Dilution** |
| **SYCP3** | Mouse | Santa Cruz Biotechnology | sc-74569 | 1:100 |
| **SYCP1** | Rabbit | Abcam | ab15090 | 1:100 |
| **MLH3** | Rabbit | Gift from B. de Massy and V. Borde | - | 1:1000 |
| Secondary antibodies: |  |  |  |  |
| **Reactivity** | **Host** | **Conjugate** | **Source** | **Dilution** |
| **Anti-mouse** | Goat | Alexa Fluor 594 | Thermo Fisher | 1:150 |
| **Anti-rabbit** | Donkey | Alexa Fluor 488 | Thermo Fisher | 1:150 |

**Supplementary Table S3.** Genotype distribution among offspring from *Fancm^+/−^;Spo16^+/−^* intercrosses.

| ***Fancm* genotype** | **Spo16<sup>+/+</sup>** | **Spo16<sup>+/−</sup>** | **Spo16<sup>−/−</sup>** | **Total** |
|---|---|---|---|---|
| ***Fancm*<sup>+/+</sup>** | 21 | 56 | 22 | **99** |
| ***Fancm*<sup>+/−</sup>** | 45 | 110 | 47 | **202** |
| ***Fancm*<sup>−/−</sup>** | 11 | 31 | 19 | **61** |
| **Total** | 77 | 197 | 88 | **362** |

